# Reduction of complex dynamic touch information to a single stable perceptual feature

**DOI:** 10.64898/2026.04.10.717744

**Authors:** Naghmeh Zamani, Benjamin Stephens-Fripp, Chase Tymms, Sonny Chan, Roham Pardakhtim, Heather Culbertson, Jess Hartcher-O’Brien

## Abstract

Dynamic touch requires the perceptual system to extract stable material properties from complex, evolving signals. Here, we examine potential material property features for a constrained dynamic-touch regime: brief taps, in which contact-induced vibratory transients are prominent and sustained deformation cues are reduced. Using a spectral-energy compensation method, we conducted five psychophysical experiments in two degraded-feedback scenarios: soft finger interfaces, in which fingertip stiffness was reduced by an inflatable silicone bubble, and compliant-surface interactions, in which participants tapped foam surfaces. Participants made explicit hardness judgments in two-alternative forced-choice tasks and identified rendered materials in multi-alternative matching and classification tasks. Under these tapping-based conditions, judgments scaled systematically with total spectral energy and were largely independent of waveform family. When spectral energy and dominant-frequency cues were placed in conflict, classifications predominantly followed spectral energy. These results identify total spectral energy as a robust cue for hardness judgments and material identification from brief taps, rather than as a complete account of hardness perception across all exploratory procedures. The findings provide a compact rendering principle for haptic interfaces in which contact-triggered vibrotactile feedback is available.

**Teaser:** During transient contacts between the skin and an object, total spectral energy is a robust cue driving hardness judgments and material identification, with less dependence on waveform family or dominant frequency.

## Introduction

When humans interact with objects through touch, they rely on material properties such as hardness, compliance, texture, and friction, which are conveyed through tactile cues including high- and low-frequency vibrations, microslips, skin strain, and slip motion to guide perception and action. A tap on a surface excites a brief transient that propagates through finger tissue as broadband vibrations. The nervous system interprets these vibrations to infer properties such as whether a surface is hard like metal or soft like foam (*1*). Hard surfaces produce higher-amplitude, higher-frequency transients, whereas soft materials yield attenuated, lower-frequency signals. Classical studies established that tactile afferent populations show broad frequency- and amplitude-dependent sensitivity, with different afferent classes responding preferentially to different ranges of vibration (*2, 3*). This physiology has motivated haptic approaches that preserve high-frequency contact transients, particularly those transmitted through Pacinian-associated channels (*4*). However, preferential sensitivity does not imply narrow-band perceptual coding: supra-threshold tactile vibrations can recruit afferents outside their preferred bands, and humans have limited ability to label precise vibration frequency, particularly at higher frequencies.

At suprathreshold levels, the total energy in a vibration signal may provide a more robust cue for certain material judgements than precise frequency labels. Neurophysiological and psychophysical studies show that afferent responses vary with vibration amplitude and power across broad frequency ranges, rather than with a single frequency alone (*5–7*). This does not imply that temporal structure is unimportant: fine timing in tactile afferent responses supports other functions, including rapid object manipulation and edge-orientation processing (*8,9*).

Natural textures and materials generate broadband vibration “signatures,” with harder materials imparting greater high-frequency energy to the skin (*1, 10, 11*). Consistent with this view, Toscani and Metzger’s ViPer database shows that perceptual judgments of natural textures are closely related to the spectral statistics of the corresponding vibratory signals (*12*). These findings suggest that the tactile system may rely on the total spectral energy - the integrated vibratory power across - rather than specific frequency components, as a compact variable for judging hardness. Similar energy-integration strategies have been observed in other sensory systems, such as vision and audition (*13, 14*).

Parallel work in robotics and artificial tactile sensing has shown that texture, hardness, and other material properties can be classified by combining diverse exploratory movements with high-dimensional features extracted from contact signals. In these systems, sliding, pressing, tapping, force trajectories, and microvibration patterns can all contribute to material classification (*15–17*). The present study has a different goal: rather than optimizing sensing or exploration for machine classification, we ask which limited vibratory information human observers can use to judge hardness and identify materials during brief taps. We therefore deliberately constrain exploration to light, contact-triggered taps, allowing the timing and energy of the rendered transient to be controlled across trials and observers.

When part of this vibratory content is lost, such as when a finger is covered by a soft interface, or when interacting with compliant or damped surfaces, perceived hardness decreases (*18–21*). Recent work with encountered-type haptic displays shows that the physical hardness of the device’s contact interface itself (e.g., a rigid plate versus a padded surface) systematically biases how stiff the same virtual object feels, consistent with the idea that attenuating high-frequency vibratory energy makes materials feel softer (*22*). Similarly, in stylus-based haptic augmented reality, contact-triggered vibrations delivered through the stylus can reliably bias judgments of how hard a surface feels (*23*). In these studies, increasing vibration amplitude and high-frequency content can be understood as increasing the spectral energy delivered to the skin. Together, these findings suggest that restoring this missing spectral energy in contact transients can recover, and even systematically alter, perceived hardness.

We operationalise perceptual effects using standard psychophysical tasks. In Experiments 1 and 3, participants made two-alternative forced-choice judgments of which contact *felt harder* ; in Experiments 2, 4, and 5, they matched or classified the rendered contact as one of several material categories. These tasks probe both a graded hardness dimension and a higher-order mapping from a contact transient to material identity; they are not vibration-detection tasks. We hypothesize that by artificially reintroducing the missing spectral energy of contact transients we can restore or alter the perception of hardness. Using this principle, one can determine the transient energy needed for a given “virtual hardness” and inject that energy at contact to trick the sense of touch. This concept shifts the focus from precise waveform replication to matching the power spectral density (PSD: a measure of how vibration intensity is distributed across different frequencies) profile that a real material would generate.

This principle is potentially relevant to prosthetics, teleoperation, and immersive virtual reality, where compliant interfaces or remote tools can attenuate natural contact transients (*4, 24–26*). Its implementation in these settings will require contact-synchronous feedback and, for richer exploratory movements, movement-dependent modulation of the rendered signal. Here, we test the more limited hypothesis that restoring controlled spectral energy at contact can systematically alter hardness judgments and material identity during brief taps. We conducted five experiments in two degraded-feedback scenarios: (i) a soft-finger interface, where fingertip stiffness was reduced, and (ii) a compliant-surface interface, where the object itself was soft (Fig. 1).

**Figure 1.**
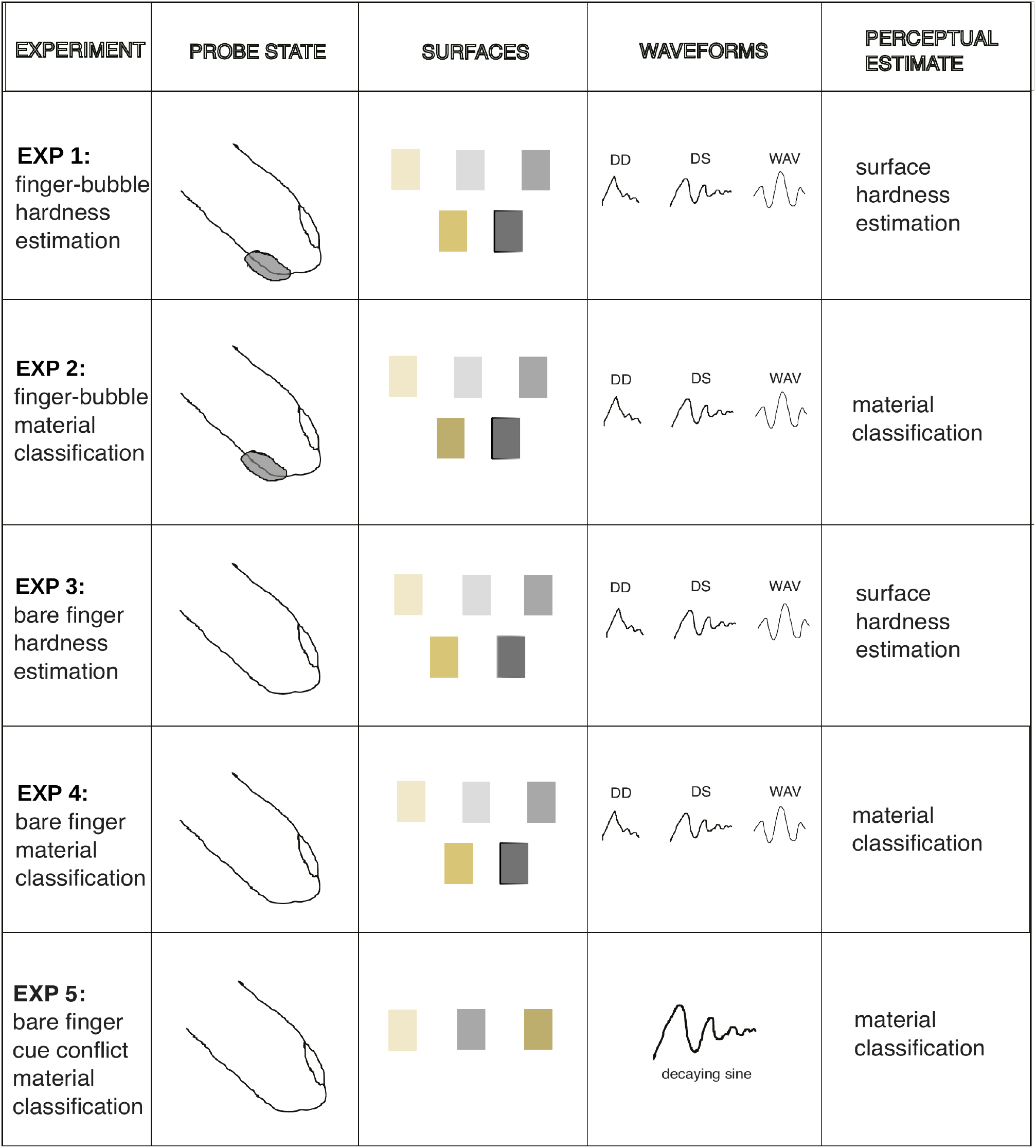
Experimental conditions across the five experiments (rows).

### Results: Spectral Energy Compensation Enables Perception of Hardness and Material Identity

We investigated whether restoring high-frequency spectral energy in contact-evoked vibrations is sufficient to recover perception of hardness and material identity when natural mechanical cues are degraded. In all experiments, participants tapped either a rigid wood block or a compliant foam block with the index finger while a nail-mounted vibrotactile actuator (Lofelt L5) replayed brief (∼50 ms) vibration bursts that were time-locked to contact via a photoelectric sensor (trigger-to-output latency *<* 5 ms; see Fig. 9). Compensation signals were constructed in the frequency domain by subtracting the spectrum of the degraded condition (soft finger or soft surface) from that of a harder reference material and then band-limiting the result to 30–1000 Hz, selectively boosting the part of the power spectral density where hard and soft contacts differed (mainly at higher frequencies). Contact onset was detected independently of impact magnitude, allowing each transient to be delivered at a consistent phase relative to the tap. The low-latency Bela implementation was selected to close the contact-feedback loop within a temporal range expected to support action-feedback synchrony.

We tested two degraded-feedback paradigms: (i) contact with a hard surface using an artificially softened fingertip, achieved by mounting an inflatable silicone bubble over the fingerpad, and (ii) contact with a soft foam surface using a bare fingertip. In both paradigms, we asked whether adding these spectrally rich transients to the degraded contact could re-enable relative hardness discrimination (2AFC) and material identification (5-AFC).

Participants experienced three families of compensation signals: (1) data-driven waveforms recorded from real taps on the reference materials, (2) decaying sinusoidal bursts tuned to each material’s dominant frequency, and (3) broadband Ricker wavelet pulses. For each family, signals were matched in total spectral energy (RMS over 30–1000 Hz; Methods, Signal Calibration) so that spectral energy, rather than waveform shape, was the primary manipulated variable. Behavioral performance was summarized with confusion matrices and analyzed using repeated-measures ANOVAs with planned post hoc comparisons (see Methods for full details).

### Compensating for Finger Softness: Restoring Hardness Perception through Spectral Energy

#### Experiment 1: Spectral vibration cues enable hardness discrimination with a soft fingertip

To simulate fingertip compliance relevant to prosthetic and wearable technologies, 10 participants wore an air-filled silicone bubble on each index finger, reducing finger stiffness, and a nail mounted vibrotactile actuator on each index finger. They tapped two identical rigid wood blocks with each index finger, where one of them randomly being delivered a vibration signal and the other silent, and judged which contact felt harder (two-alternative forced choice (2AFC) task). Vibration total spectral energy varied across five discrete levels corresponding to the five different materials, each delivered using one of three waveform types (data-driven, decaying sinusoid, or Ricker wavelet).

Participants reliably judged contacts accompanied by higher spectral energy as harder (Fig. 2). Performance was near ceiling for most comparisons: Metal was judged harder than No signal, Foam, Silicone, and Rubber on 100% of trials, and harder than Wood on 85-90% of trials (depending on waveform). Similarly, Wood was consistently judged harder than all softer materials (100%). The only systematic confusions arose at the softest contrast: Silicone was judged harder than Foam on only 20-25% of trials (*p >* 0.99, binomial test). Rubber was judged harder than Foam on 75-85% of trials (*p <* 0.01), and reliably harder than Silicone and No signal (*p* ≪ 0.001). These results establish that participants’ hardness judgments were overwhelmingly consistent with spectral energy differences, except at the smallest increment (Foam-Silicone).

**Figure 2.**
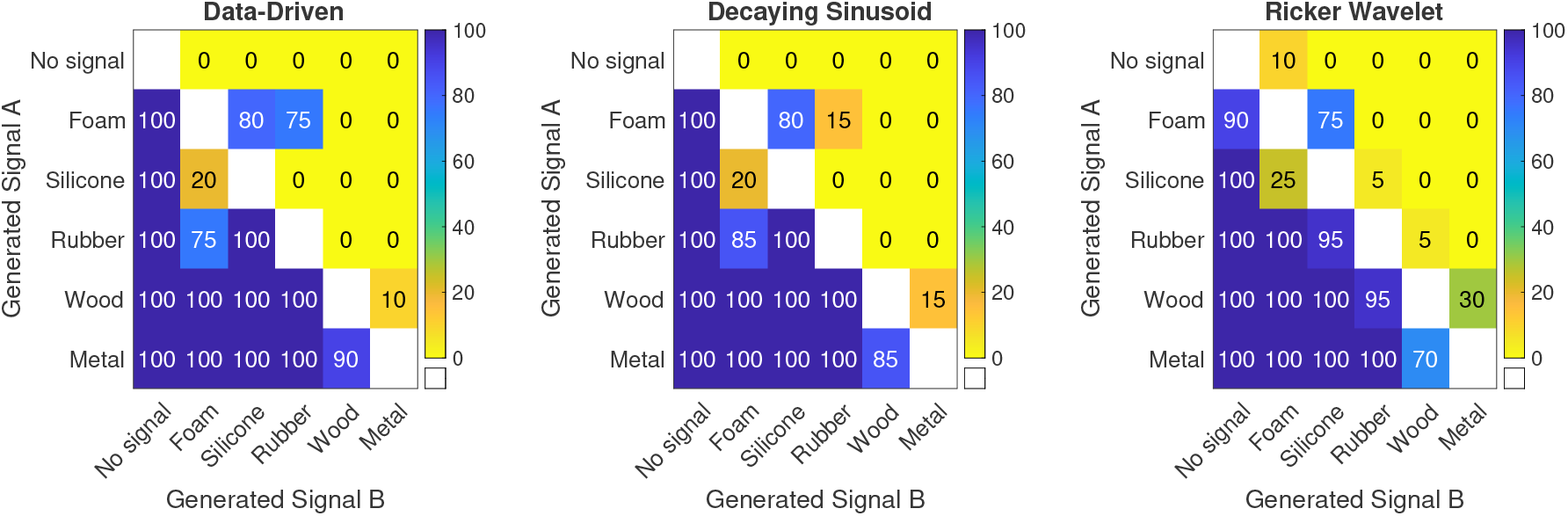
Hardness discrimination with softened fingertips. Participants wore an inflated bubble to reduce fingertip stiffness and tapped two identical rigid blocks; one tap was augmented with a brief vibration. Confusion values denote the proportion that the row stimulus was judged harder than the column stimulus. Judgments increase monotonically with the delivered total spectral energy, largely independent of waveform family.

Binomial tests on the four adjacent contrasts confirmed this pattern. Foam-Silicone performance did not exceed chance (20-25%, *p >* 0.99), whereas Silicone-Rubber and Rubber-Wood were discriminated at or near ceiling (≥95%, all *p <* 10^*−*6^). Wood-Metal performance was strong but less than perfect: 90% (Data-driven, *p <* 10^*−*5^), 85% (Decaying Sine, *p <* 10^*−*4^), and 70% (Wavelet, *p* = 0.012).

A two-way repeated-measures ANOVA on per-subject accuracies confirmed a robust main effect of difficulty (*F* (3, 27) = 22.49, *p* = 1.65 *×* 10^*−*7^, 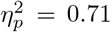), reflecting the progressive increase in accuracy from Foam-Silicone to Wood-Metal. No main effect of waveform was observed (*F* (2, 18) = 1.91, *p* = 0.18), nor any interaction (*F* (6, 54) = 1.63, *p* = 0.16), indicating that all waveform types yielded the same difficulty-dependent pattern. One-way ANOVAs within each waveform similarly showed strong effects of difficulty (all *p <* 0.001). Post hoc paired comparisons confirmed that Foam-Silicone was significantly worse than all other contrasts (Holm-corrected *p <* 0.01).

To assess global discriminability, we pooled trials where each nominal hardness level was paired against all softer levels. A two-way repeated-measures ANOVA (hardness level x waveform) revealed a strong main effect of hardness level (*F* (4, 36) = 78.52, *p* = 3.1 × 10^*−*15^, 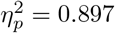), but no effect of waveform (*F* (2, 18) = 1.84, *p* = 0.19) and no interaction (*F* (8, 72) = 1.02, *p* = 0.43). Binomial tests showed ceiling-level discrimination for Metal and Wood against all softer levels (all *p* ≪ 0.001), high performance for Rubber (≥95%), and near-chance performance only for Foam-Silicone. This confirms that hardness perception is preserved across all but the smallest spectral energy increment.

Together, these results demonstrate that high-frequency spectral energy is sufficient to restore hardness perception through a softened fingertip. The somatosensory system appears to prioritize total energy content over waveform details when interpreting tactile transients, allowing reliable discrimination except when energy differences are minimal.

#### Experiment 2: Synthetic vibration cues support material identity with a soft finger

We next asked whether spectral vibration cues could not only restore perceived hardness but also deliver material-specific qualities. 10 participants tapped a rigid wooden block with a compensation vibration signal, then explored five reference materials, foam, silicone, rubber, wood, and metal - using their unaltered index finger. The materials increased in stiffness from foam (*E* ≈0.01 GPa) to metal (*E* ≈200 GPa). The task was to match the augmented wood tap to the closest material.

Participants’ choices aligned strongly with spectral energy (Fig. 3). The no-vibration condition was almost always matched to foam, intermediate signals to silicone or rubber, and the strongest signals to wood or metal.

**Figure 3.**
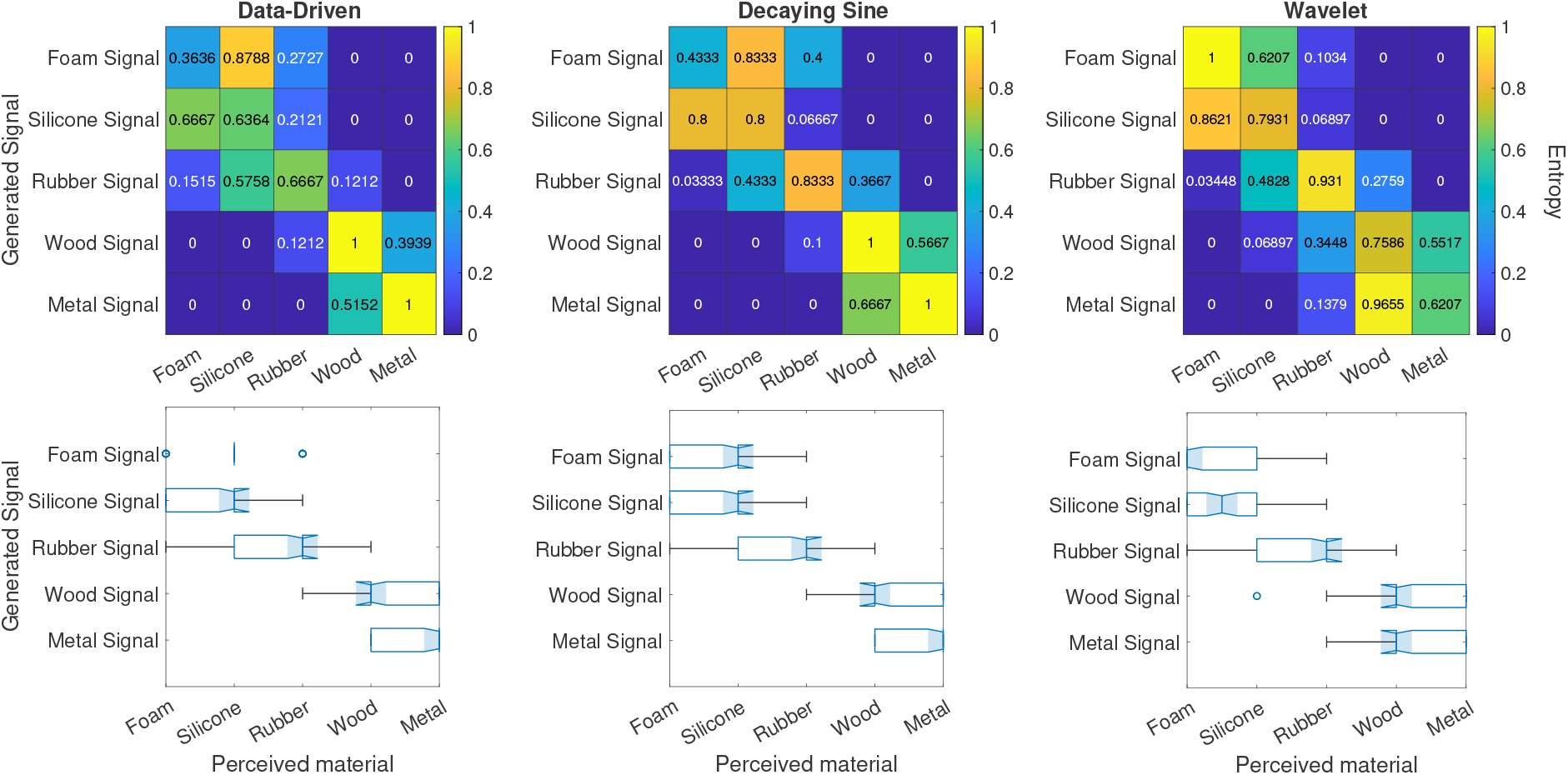
Material identification with a soft fingertip. A wood tap, felt through a bubble-softened fingertip, augmented by synthetic vibration was matched to one of five real materials. Response distributions shift systematically from foam→metal as total spectral energy increases, indicating that energy conveys material identity even when fingertip mechanics are softened.

Confusion matrices confirmed a near-diagonal structure with only nearest-neighbor errors at the soft and mid-levels. Overall, 67% of responses were exact matches, and 95% were within ±1 material rank of the intended target (*p <* 0.001, binomial), indicating highly systematic scaling. Ordinal correlation between signal energy and perceived material rank was strong across all waveforms (*r* = 0.82 − 0.84). Mutual information analysis showed that the vibration cues carried 0.63-0.69 bits of categorical information (out of a theoretical maximum of 2.32 bits), indicating substantial transmission of material identity through synthetic vibrations.

A two-way ANOVA (signal level x waveform) on response ranks revealed a robust main effect of signal level (*F* (4, 749) = 177.23, *p <* 0.001, *η*^2^ = 0.76), no main effect of waveform (*F* (2, 749) = 2.34, *p* = 0.158), and a minor but significant interaction (*F* (8, 749) = 3.55, *p <* 0.001, *η*^2^ = 0.009). Tukey post hoc tests confirmed that each increase in vibration level produced significantly harder perceived materials (all adjacent contrasts *p <* 0.01), with the exception that the two highest levels (wood vs. metal signals) were not reliably different (*p* = 0.12). This indicates that perceptual scaling was systematic but compressed at the upper end of the stiffness range.

These findings indicate that synthetic vibration cues not only restored a sense of hardness but also conveyed material-specific identity. Even when fingertip mechanics were altered to simulate soft robotic contact, vibration energy was sufficient to drive perceptual categorization along the natural continuum of material stiffness.

### Compensating for Soft Surfaces: Vibration Signals Restore Hardness Perception

#### Experiment 3: Graded hardness perception on a soft surface via spectral vibration cues

Here, we reversed the scenario: 10 participants tapped two identical soft foam blocks using bare fingertips with nail-mounted vibrotactile actuators, where one of which delivered a vibration burst (2AFC). Five energy levels and three waveform types (Data-driven, Decaying Sine, Wavelet) were tested.

Participants reliably judged contacts accompanied by higher spectral energy as harder (Fig. 4). Performance was near ceiling for most comparisons: Metal was judged harder than No signal, Silicone, and Rubber on 100% of trials, and harder than Wood on 75-80% of trials depending on waveform. Wood was consistently judged harder than No signal and Silicone (100%), and harder than Rubber on 85-95% of trials. Rubber was judged harder than No signal and Silicone on 95-100% of trials, and Silicone was always judged harder than No signal. These results establish that hardness judgments closely tracked spectral energy differences across the foam substrate, with the only systematic errors arising in the Wood-Metal contrast.

**Figure 4.**
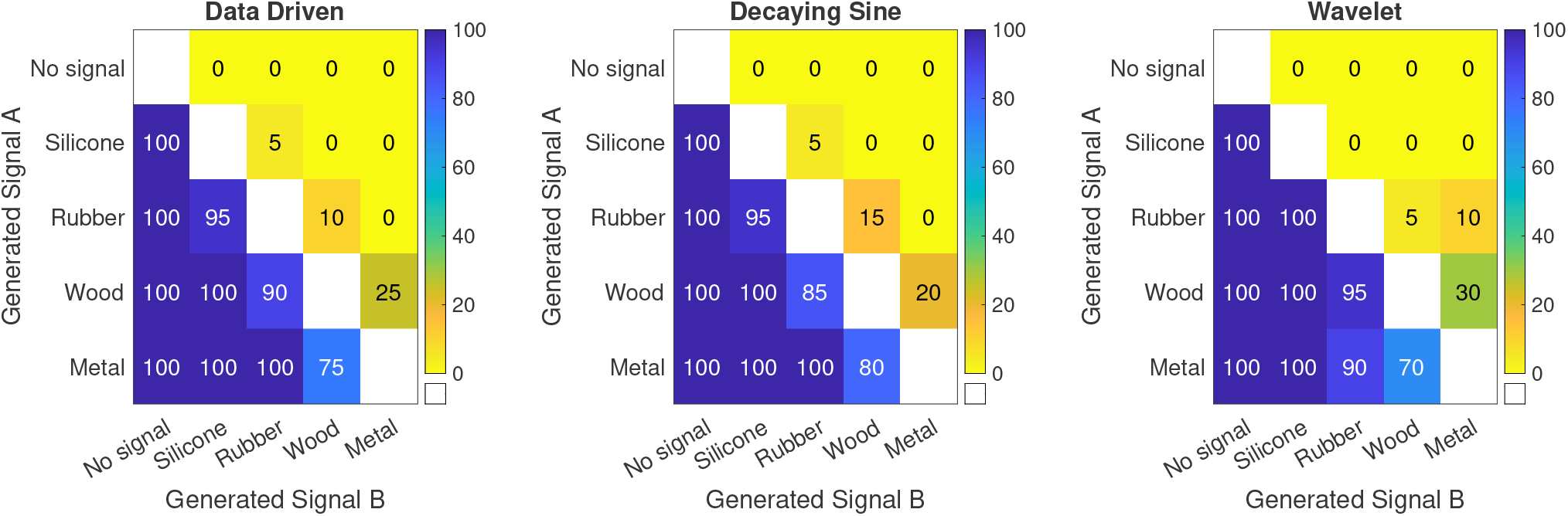
Hardness discrimination on a compliant surface. With bare fingers, participants tapped identical foam blocks; one tap included a vibration burst. Confusion matrices show that perceived hardness follows the signal’s total spectral energy across waveform types, with small residual confusions only at the highest two levels (wood-metal).

Binomial tests on the four adjacent contrasts confirmed this pattern. No signal-Silicone performance was at ceiling (100%, *p* = 8.3*×*10^*−*25^ across all waveforms). Silicone-Rubber and Rubber-Wood were discriminated at 92.5-100% (*p* ≤ 2.7 *×* 10^*−*21^). Wood-Metal performance was strong but less than perfect: 87.5% (Data-driven, *p* = 1.6 *×* 10^*−*12^), 90% (Decaying Sine, *p* = 2.7 *×* 10^*−*14^), and 85% (Wavelet, *p* = 6.0 *×* 10^*−*11^).

A two-way repeated-measures ANOVA on per-subject accuracies showed a trend-level main effect of difficulty (*F* (3, 27) = 2.92, *p* = 0.052, 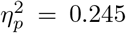), no main effect of waveform (*F* (2, 18) = 0.11, *p* = 0.893), and no interaction (*F* (6, 54) = 1.83, *p* = 0.111). One-way ANOVAs within each waveform showed a significant effect of difficulty for Wavelet (*F* (3, 27) = 3.55, *p* = 0.028), and trend-level effects for Data-driven (*F* (3, 27) = 2.47, *p* = 0.083) and Decaying Sine (*F* (3, 27) = 2.00, *p* = 0.138). Thus, although the across-subject variability reduced the omnibus effect, all contrasts were significantly above chance, though Wood-Metal fell below ceiling.

To assess global discriminability in Experiment 3, we again pooled trials where each nominal hardness level was paired against all softer levels. The omnibus ANOVA showed only a trend-level effect of hardness level (*F* (3, 27) = 2.72, *p* = 0.064), with no main effect of waveform or interaction (*F* (2, 18) = 0.046, *p* = 0.955; *F* (6, 54) = 1.87, *p* = 0.102). However, pooled binomial tests corroborated the monotonic scaling, with ceiling performance for Silicone (100%, *p* = 5.66 *×* 10^*−*73^) and near-ceiling for Rubber (99.2%, *p* = 7.06 *×* 10^*−*136^), Wood (98.3%, *p* = 6.81 *×* 10^*−*192^), and Metal (96.5%, *p* = 4.99 *×* 10^*−*227^).

Together, these results mirror Experiment 1, demonstrating that the somatosensory system prioritizes vibratory spectral energy over substrate stiffness when inferring hardness. Even on a uniformly soft foam surface, participants’ judgments followed the delivered signal’s spectral energy, confirming that tactile transients can override mechanical compliance cues.

#### Experiment 4: Material identification on a soft substrate via synthetic vibration

In Experiment 2, a bubble-softened finger tapped a rigid wood block and spectral vibration cues conveyed material identity with high fidelity. Here we asked whether the same cues remain effective when the fingertip is bare and the surface is compliant foam. 10 participants tapped the foam block, received one of four vibration signals, and matched the percept to one of five reference materials.

Perceived material identity continued to scale with vibration intensity (Fig. 5). Low-energy signals were judged as foam or silicone, mid-levels as silicone or rubber, and high-energy signals as wood or metal. Confusion matrices confirmed a strong diagonal structure, with errors predominantly restricted to nearest-neighbor categories. Overall, 52% of responses were exact matches and 94% were within ±1 material rank of the intended target (*p <* 0.001). Ordinal correlations between signal level and perceived material rank remained significant across all waveform types (*r* = 0.62 − 0.65), though weaker than in Experiment 2 (*r* ≈0.83). Mutual information analysis showed that the vibration cues carried 0.29-0.31 bits of categorical information, roughly half of the information transmitted in Experiment 2, indicating reduced precision relative to Experiment 2.

**Figure 5.**
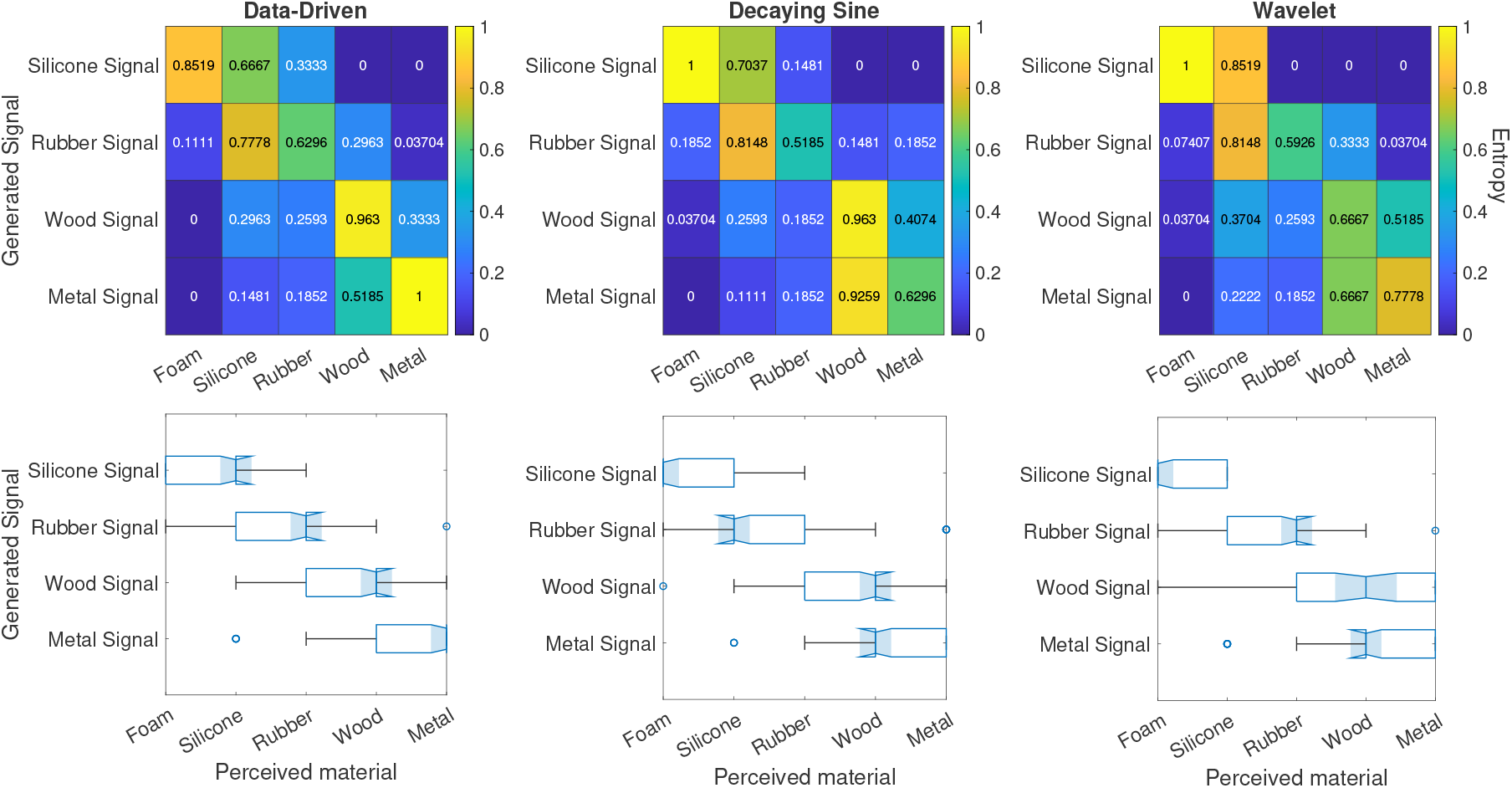
Material identification on a soft substrate (soft foam surface; bare fingertip). Foam taps augmented by synthetic vibrations were matched to real materials. Despite reduced precision relative to Experiment 2, perceived category still scales with total spectral energy, shifting from foam/silicone at low levels to wood/metal at high levels.

A two-way ANOVA (signal level x waveform) confirmed a robust main effect of signal level (*F* (3, 599) = 642.01, *p <* 0.001, *η*^2^ = 0.55), but no effect of waveform type (*F* (2, 599) = 2.18, *p* = 0.194) and no interaction (*F* (6, 599) = 0.38, *p* = 0.89). Tukey post hoc tests showed that each increase in signal level produced a significant shift toward harder perceived materials (all adjacent contrasts *p <* 0.01), except for the highest comparison (wood vs. metal), which was not consistently discriminated (*p* = 0.21). This confirms that perceptual scaling remained systematic but compressed at the upper end of the stiffness range.

Together, these results demonstrate that, although categorical precision was lower than in Experiment 2, spectral vibration cues continue to systematically drive material identity judgments.

#### Experiment 5: Perceived Material Identity Is Dominated by Rendered Spectral Energy

To directly test whether the somatosensory system prioritizes total spectral energy over frequency when inferring material properties, we designed stimuli for which these cues were intentionally put in conflict, e.g., combining the RMS amplitude of foam with the frequency characteristics of wood. This allowed us to dissociate the contribution of each cue and determine the behaviorally relevant material feature. We conducted a material classification task with a reduced stimulus set (foam, rubber, wood) comprised of such cue conflict stimuli. On each trial, participants tapped a physical foam block with a bare finger while receiving a vibration signal - corresponding to the RMS of one of the three materials and a base frequency of one of the other materials - delivered to the dorsal side of the fingertip. This setup created congruent and incongruent combinations of base and rendered cues.

Participants classified the perceived material from among the three options. Confusion matrices (aggregated across 10 participants, five repetitions per condition) revealed that judgments overwhelmingly aligned with the rendered vibration rather than the actual material of contact (Fig. 6).

**Figure 6.**
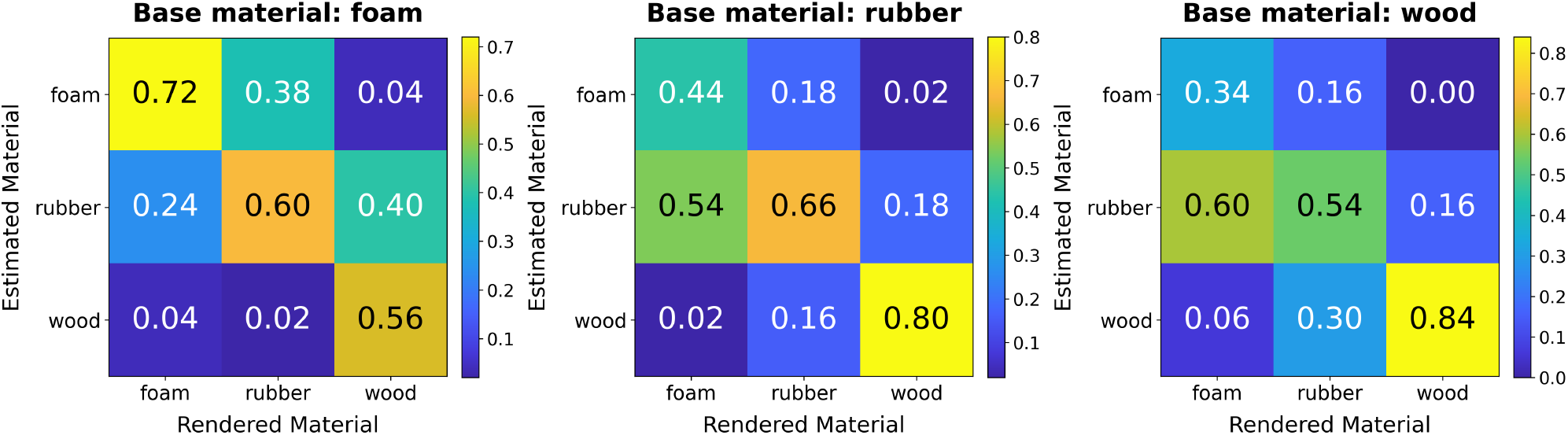
Cue-conflict classification. While tapping foam, participants received brief vibration transients in which the dominant frequency class and the RMS (total spectral energy) were crossed. i.e the transient was constructed to use the dominant frequency of the base material, but containing the total spectral energy of the rendered material. Choices tracked the rendered spectral energy (RMS target) rather than dominant frequency or base material, demonstrating that total spectral energy dominates perceived material identity under conflict.

**Figure 7.**
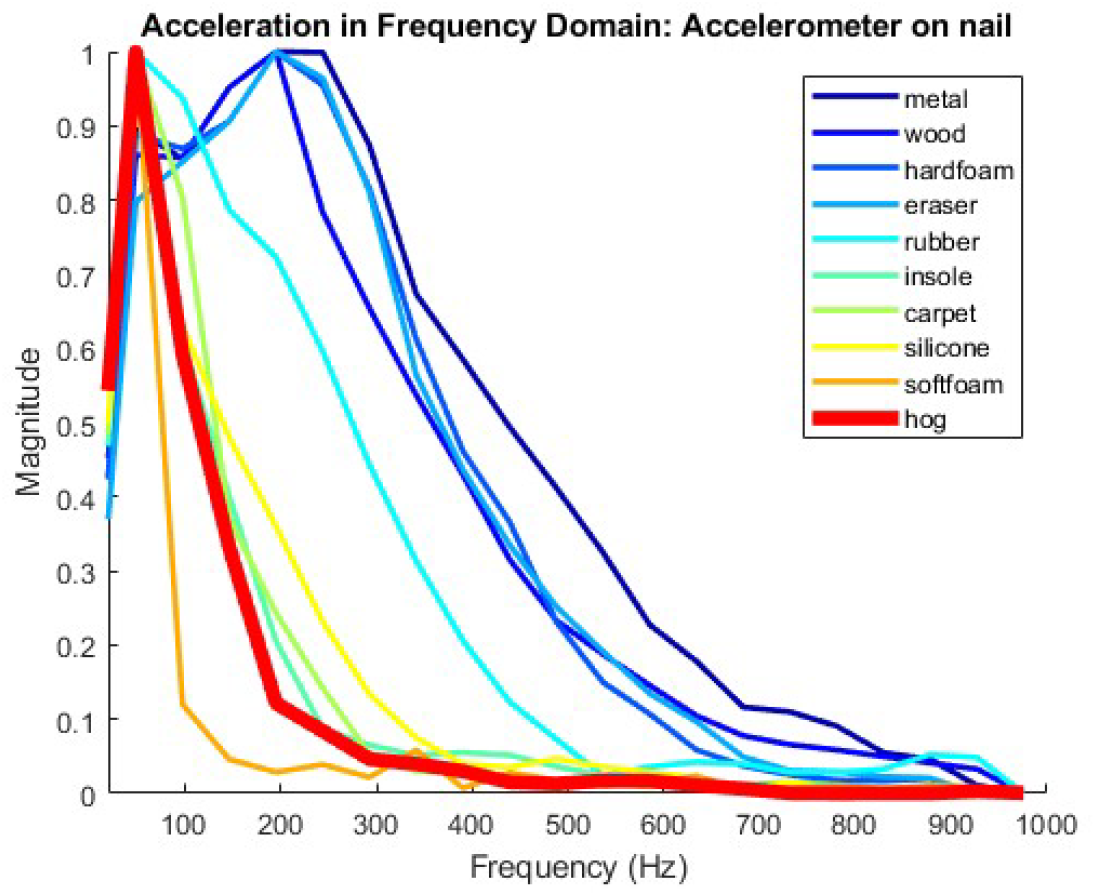
Spectral analysis of vibration recordings from 20 everyday materials. The clustering was used to select repre-sentative materials spanning the recorded spectral-response space for the main experiments.

**Figure 8.**
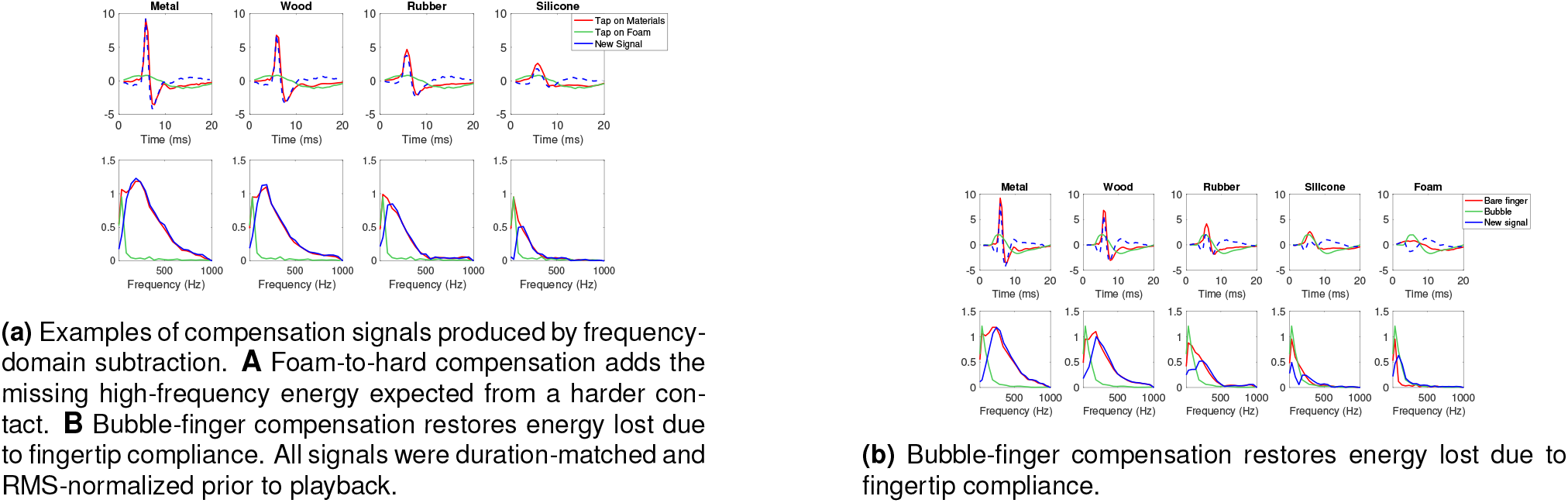
Examples of processed compensation signals generated via frequency-domain subtraction.

A binomial logistic regression confirmed a significant effect of the rendered signal. In particular, the “wood” vibration strongly influenced judgments (*β* = 1.015, *p <* 0.001; odds ratio = 2.76, 95% CI [1.70, 4.48]), while “rubber” showed a positive trend (*β* = 0.407, *p* = 0.082). By contrast, the physical base material had no significant effect.

Model comparison further supported this interpretation. Including interaction terms between base frequency and RMS rendered materials significantly improved fit (AIC = 572.99 vs. 592.41; likelihood ratio test *χ*^2^(4) = 27.42, *p <* 0.001). Nonetheless, the rendered spectral energy RMS component remained the dominant determinant of perceived material identity.

These results confirm that, when presented with conflicting cues, the tactile system relies more heavily on the spectral characteristics of vibratory input than on the fundamental frequency response of the contacted surface. Total spectral energy - not frequency-specific patterns or physical compliance - drives material classification under ambiguity.

## Discussion

Across five experiments, we show that total spectral energy is a robust cue for graded hardness judgments and material identification during brief tapping interactions. The effect was observed in both soft-finger and compliant-surface paradigms. Under energy-matched conditions, simple synthetic waveforms, including decaying sinusoids and wavelets, produced response patterns broadly similar to recorded contact transients. In Experiment 5, when spectral energy and dominant-frequency cues were placed in conflict within the vibrotactile signal, material classifications predominantly followed spectral energy.

These findings identify a compact cue for a specific dynamic-touch regime, rather than a universal organizing principle for all haptic perception. Our experiments deliberately emphasized brief contact transients and reduced the availability of sustained deformation, force, and proprioceptive signals. Consequently, the cue conflict in Experiment 5 was within the vibrotactile domain; it does not establish how spectral energy will be weighted against strong non-vibratory or cross-modal cues. In other sensory systems, cue combination can reflect reliability-weighted averaging, context-dependent dominance, or dynamic reweighting. Determining which of these regimes governs tactile hardness will require experiments that place spectral energy in conflict with sustained force, skin stretch, proprioception, and audition.

Our findings demonstrate that **matching the spectral energy of contact vibrations** can robustly convey perceptions of material hardness, even under conditions where mechanical cues are degraded or misleading. Across five experiments, we show that participants reliably perceived gradations in hardness - and, in many cases, could identify specific materials - based solely on vibration cues, despite interacting with physically soft or compliant surfaces.

Crucially, this perceptual reconstruction occurred regardless of the waveform’s temporal structure - whether it was a natural recording, a sinusoidal burst, or a wavelet pulse - as long as the spectral energy matched the corresponding hardness level. These results provide strong evidence that the human tactile system prioritizes

*spectral energy* over precise waveform features as a core cue in inferring hardness.

These insights extend current models of tactile perception. Classical frameworks emphasize force-displacement dynamics or sustained pressure, typically mediated by slowly adapting mechanoreceptors, as primary cues for stiffness and compliance (*27*). However, in dynamic touch contexts such as tapping, brief high-frequency transients dominate the tactile signal (*6*), suggesting a distinct perceptual mechanism. Our results support the hypothesis that **dynamic spectral energy - particularly within the 100–300 Hz band - serves as a proxy for mechanical hardness**, interpreted by the nervous system even in the absence of congruent proprioceptive or mechanical feedback.

One of the most striking findings was the perceptual reconstruction exhibited by participants: despite interacting with a physically softer interface, brief high-frequency vibrations were sufficient to evoke the illusion of a hard material. This result shows that, in the constrained tapping conditions tested here, brief high-frequency vibrations can substantially cue the perceptual reconstruction of perceived hardness despite a physically compliant contact interface. It does not establish that spectral energy overrides proprioceptive or mechanical cues generally, because those cues were not independently manipulated or placed in controlled conflict.

From an engineering standpoint, this perceptual flexibility presents a valuable design opportunity. Rather than relying on complex force-feedback systems - which are often unstable, costly, or difficult to miniaturize - haptic devices can instead leverage **computationally efficient, precomputed vibration signals** to simulate hardness percepts. The Lofelt L5 was chosen for experimental control rather than wearable compactness. Because waveform family had relatively little effect under energy-matched conditions, the approach should be compatible with smaller broadband transducers, especially piezoelectric actuators, which are well suited to reproducing the high-frequency content relevant to distinctions such as wood versus metal, provided that they can deliver the required energy without relying on a narrow resonant response. In tandem with existing physics engines, rendering can additionally leverage the spectral energy cues that would be computed as a function of a finger interacting with a given material.

We masked audible tapping sounds to isolate vibrotactile contributions and to reduce sound-biased touch effects, including the parchment-skin illusion (*28*). Thus, the observed energy-dependent hardness judgments and material classifications did not require auditory cues. Precisely timed sounds could nevertheless provide an alternative or complementary rendering channel, a question not tested here. More generally, cross-modal substitutions may differ in uncertainty and robustness across environments and observers; mechanically coupled vibration transmitted through the skin may offer a comparatively direct and noise-resistant route for rendering contact transients. Testing audio-only and audio-plus-vibration rendering is an important next step.

Importantly, the perceptual fidelity required for realism appears to depend more on **accurate spectral energy matching and precise temporal alignment with contact onset**, rather than on detailed modeling of mechanical interactions or force dynamics (*29, 30*). Timing of these haptic feedback cues is critical given the millisecond encoding precision of AC-coupled Pacinian corpuscles, tuned to transient impacts and their spectral content (*31, 32*). Accounting for these dynamics dramatically increases skin sensing capabilities, as Rongala et al. demonstrate in their neural simulation work (*32*). Follow-up investigations aimed at quantifying the perceptual limits of these timing constraints could establish concrete design tolerances for haptic systems. Identifying the temporal resolution required for reliable material discrimination would not only refine rendering strategies, but also inform hardware specifications for next-generation tactile interfaces. Future work characterizing the perceptual bounds of these temporal dynamics would provide principled constraints for tactile display design. By mapping the thresholds at which temporal imprecision degrades material percepts, such studies could bridge neural encoding models and practical engineering implementations.

Our tasks probe two linked perceptual outcomes. In the 2AFC experiments, total spectral energy systematically shifted a graded hardness judgment: which contact felt harder. In the matching and classification experiments, observers mapped the rendered transient onto learned-over-the-lifespan material categories. The latter judgment is necessarily more categorical and may be influenced by additional spectral and contextual information. Thus, our data show that total spectral energy is sufficient to organize both hardness judgments and material-category responses in the tested tapping regime, without implying that it exhausts the information used for material identification during unconstrained touch.

These findings do not support a simple account in which hardness judgments from brief taps are determined by a precise frequency label or by a cochlear-like bank of narrow frequency filters. Instead, in the dynamic-touch regime studied here, the relevant computation appears to be a task-dependent integration of spectral energy. This conclusion is compatible with evidence that fine temporal and spatial patterning in tactile afferent activity remains essential for other tasks, including object manipulation and edge-orientation processing (*8, 9*).

In practical terms, this leads to several design implications:

- **Latency is critical**. In all experiments, vibrations were triggered within ∼5 ms of contact, aligning temporally with the skin’s mechanical response. Delays could degrade the realism or effectiveness of the illusion.
- **Calibration is tractable**. We demonstrated that participants can reliably map a vibration signal to a material, implying that users could calibrate their own haptic devices via interactive matching.
- **Personalization may help**. Participants differed slightly in sensitivity to vibration levels, suggesting future systems could learn user-specific thresholds or preferences.

Of course, our approach has limitations. It focuses on single, transient taps rather than sustained pressure, indentation, or squeezing. A particularly informative next experiment would compare materials matched in static modulus but differing in damping or dynamic stiffness, tested with bare-finger taps. Such a manipulation would dissociate static stiffness from the vibration signatures elicited by impact and determine more directly how total spectral energy is weighted relative to other mechanical cues. In real-world manipulation, hardness perception is multimodal and dynamic, so **vibratory cues should be seen as augmentative**, not comprehensive to rendering the interaction. However, that is precisely the nature of a perceptual feature - a signature used by the sensory system to successfully reconstruct a property of the environment. In applications such as **teleoperation, prosthetics**, or **VR/AR interfaces** - comprehensive mechanical feedback is unavailable. The present results should therefore be interpreted as applying to tapping-based, vibration-dominated interactions, not as a complete account of hardness perception during all natural exploratory procedures.

In summary, this study introduces and validates a **spectral energy compensation framework** for rendering material hardness through high-frequency contact transients. By controlling the total spectral power, regardless of waveform, we were able to restore or simulate tactile hardness across different degraded interaction conditions. These findings establish high-frequency spectral energy as a reliable proxy for stiffness, supporting both graded hardness discrimination and material identification in the absence of realistic force cues.

This principle offers a path forward for lightweight, low-power haptic systems that rely on transient overlays instead of complex actuators. By aligning artificial transients with the perceptual mechanisms of the skin, haptic experiences can be made both intuitive and adaptable to diverse applications, from **robotic touch** to **prosthetic feedback** to **immersive virtual environments**.

## Materials and Methods

### Overview

We conducted five experiments to evaluate how spectral energy cues in vibration signals contribute to material perception, particularly in the encoding of hardness via touch. These experiments manipulated either *fingertip compliance* or *surface properties* and employed three signal-generation techniques. All experiments used controlled tapping procedures, precise haptic playback, and randomized trial designs.

### Participants

#### Participants (Experiments 1–4)

Forty adults (22–52 years; 13 female, 27 male; 39 right-handed, 1 left-handed) were recruited. Ten participants took part in each of Experiments 1–4. All reported normal tactile sensitivity and no history of neurological or musculoskeletal disorders affecting the hands. Prior experience with haptics was self-reported as: none (2), limited (16), moderate (13), extensive (9).

#### Participants (Experiment 5)

A separate cohort of ten adults (28–51 years; 4 female, 6 male; 9 right-handed, 1 left-handed) completed Experiment 5. All reported normal tactile sensitivity and no history of neurological or musculoskeletal disorders affecting the hands.

All participants for Experiments 1-5 provided written informed consent and all procedures were approved by the Reality Labs Research ethics committee under the AGOI protocol.

### Apparatus

Vibrations were delivered via Lofelt L5 actuators affixed to the fingernail. We selected the L5 because its broad-band, non-resonant voice-coil response supported controlled rendering of brief transients over the 30–1000 Hz band and enabled spectral energy to be varied independently of waveform family. Stimuli were triggered by photoelectric sensors and played back using a Bela real-time platform with measured trigger-to-output latency below 5-ms. We used Bela rather than a conventional data-acquisition playback path because its low-latency architecture allowed feedback to be tightly time-locked to contact onset. Fingertip compliance was manipulated using an air-filled silicone bubble inflated to 5 psi and secured over the index fingerpad. Participants were blindfolded during tasks to eliminate visual cues. To minimize auditory contributions, participants wore over-ear headphones playing continuous pink noise at approximately 65–70 dB SPL throughout all tasks. Although the L5 is relatively large for multi-finger wearables, the spectral-energy compensation principle is not specific to this actuator: compact broadband piezoelectric actuators or non-resonant voice-coil actuators with sufficient output in the relevant band could implement the same approach.

### Material Selection and Vibration Recording

We created a common library of vibration transients from taps on everyday materials using a tri-axial accelerometer affixed to the fingernail. These recordings were obtained before the psychophysical studies and were used to generate the stimulus set presented to every participant; we did not record participant-specific material signals. Each material was tapped five times at a metronome-controlled pace (90 BPM) by the same experimenter at a consistent height of fully raised pointer finger without lifting hand. We selected the transient with median peak acceleration as the representative recording for that material. This common recording library was processed to generate all stimuli used across participants; participants did not generate their own recorded stimuli.

### Signal Generation

Three signal types were tested: 1. **Data-driven:** recorded transients replayed directly. 2. **Decaying sinusoid:** exponentially decaying bursts capturing resonant modes. 3. **Ricker wavelet:** broadband pulses simulating impact transients. All signals were RMS-normalized and matched in duration. Equations and parameter values are provided in Supplementary Methods.

### Experimental Procedures

Participants completed five experiments: - **Exp. 1: Soft finger discrimination** - judged which of two wood contacts felt harder (2AFC). - **Exp. 2: Soft finger matching** - matched augmented wood taps to real materials. - **Exp. 3: Soft surface discrimination** - judged hardness between two foam contacts. - **Exp. 4: Soft surface matching** - matched augmented foam taps to real materials. - **Exp. 5: Cue conflict classification** - judged material identity when spectral energy and frequency cues conflicted. Trial counts, randomization, and timing are detailed in Supplementary Methods.

### Signal Calibration

Stimuli were RMS-normalized across methods to ensure that differences in perception reflected spectral energy, not raw amplitude. A pilot study confirmed matched perceptual intensity.

### User Studies

Participants completed five experiments grouped into two categories: fingertip modification (Experiments 1-2) and surface modification (Experiments 3-5). All tasks involved either discrimination (2AFC) or material matching, with randomized trial sequences and practice sessions. Fingers were wrapped with tape to minimize confounding cues, and participants were blindfolded to eliminate visual input. At the start of each experiment participants were given guidance on the speed and height of tapping in an attempt to keep the speed and force of impact relatively consistent.

#### Experiments 1-2 (Fingertip modification)

To simulate reduced fingertip stiffness, participants wore an inflatable bubble actuator on the index finger. In Experiment 1, they tapped identical wood blocks (one with vibration playback) and judged which felt harder. In Experiment 2, they tapped a wood block with vibration and matched it to one of five real reference materials (foam, silicone, rubber, wood, metal).

#### Experiments 3-4 (Surface modification)

Here, participants tapped soft foam surfaces without fingertip modification. Experiment 3 used a discrimination task (2AFC), while Experiment 4 used the same material-matching task as in Experiment 2.

#### Experiment 5 (Cue conflict)

A separate cohort of 10 participants completed a classification task in which spectral energy (RMS) and base frequency cues were dissociated. Participants tapped foam with synthetic vibrations designed to conflict in energy vs. frequency signatures, and judged which material was perceived.

Across all studies, vibration signals were delivered within 1 ms of contact, and task structure ensured single-blind conditions. Full trial counts, durations, and detailed procedures are reported in the Supplementary Methods.

## Acknowledgments

This work was funded by Reality Labs Research. The authors declare no competing interests.

## Supplementary Methods

### Apparatus Details

Vibration feedback was delivered using Lofelt L5 actuators (30–1000 Hz) affixed to the fingernail of each index finger using double-sided tape and wax. Actuator placement was informed by Birznieks et al. research that demonstrated that mechanoreceptors around the fingernail borders reliably respond to forces applied to the finger pad (*27*). A Bela board running a real-time Linux OS (Xenomai) ensured low-latency playback (*<* 1 ms). Finger presence was detected via Panasonic CX-400 photoelectric sensors positioned directly above the contact surface. Tactile stimuli were played back using one of three signal-generation methods (described below). The soft bubble actuator used to manipulate fingertip compliance was manually inflated to and maintained at 5 psi throughout the experiment. The bubble was secured with elastic tape to the fingerpad of the participant’s index fingertip. Participants were blindfolded during all tasks and wore lightly wrapped tape on their index fingers to minimize non-vibratory tactile cues.

### Material Selection and Frequency-Domain Clustering

#### Vibration Recording

To characterize the vibratory responses of everyday materials, we recorded transients from taps on 20 samples, including natural and synthetic surfaces (see Table S1). Vibrations were measured using a tri-axial piezoelectric accelerometer (Endevco Model 35A, Endevco Corp., NY, USA; sensitivity 10 mV/g, resonance frequency ¿30 kHz), affixed to the participant’s index fingernail with orthodontic wax to ensure stable coupling. Signals were conditioned using an Endevco 133 signal conditioner and digitized at 2000 Hz via a National Instruments NI-9205 data acquisition module. Each material was tapped five times at a controlled pace (90 BPM metronome), and signals were stored for subsequent analysis and playback.

#### Fingertip Compliance Interface (Bubble Actuator)

To simulate changes in fingertip stiffness, we fabricated a soft inflatable “bubble” actuator from thermoplastic elastomer. The actuator was secured over the index fingerpad with elastic cloth tape and inflated to 5 psi using a hand pump with inline pressure gauge. This setup reduced effective fingertip stiffness while leaving the fingernail accessible for actuator mounting. Although vibration recordings were collected for all materials with the bubble attached, only the bubble-on-wood condition was used in the experiments. Wood was selected as a mid-range reference material to simplify comparisons across signal types.

#### Material Grouping and Selection

Spectral analysis was used to identify representative materials for the main experiments. For each sample, the FFT magnitude spectrum was computed, and spectral energy was summed into 50 Hz bands from 0–1000 Hz. Materials were grouped using *k*-means clustering (*k* = 4) based on spectral centroid and broadband energy. The pilot perceptual grouping exercise was used only to select a material set spanning a range of perceptual and spectral responses; it was not a preregistered or quantitative precursor of Experiment 1. Blindfolded participants were allowed to explore the materials more freely than in the main experiments. Under this richer and less constrained exploration, hard materials (e.g., metal, wood, glass, and rock) tended to be grouped as similar, whereas the softer materials were more readily differentiated. In Experiment 1, by contrast, exploration was restricted to brief light taps through a softened fingertip, and the delivered vibration was manipulated parametrically. Thus, the pilot and Experiment 1 differ in both exploratory procedure and available cue set; they were not expected to yield identical perceptual-similarity structures.

### Supplementary Methods: Signal Processing and Implementation

#### Signal Extraction and Windowing

Raw vibration recordings were collected from taps on each material at a metronome-controlled pace (90 BPM) to ensure consistent impact velocity. For each material, the tap with median peak acceleration was selected for further processing. A 20 ms analysis window (5 ms before peak to 15 ms after peak) was extracted to capture the transient response. To reduce onset artifacts and spectral leakage, the signal was zero-padded and tapered with a 5 ms Hanning window.

#### Compensation Signal Design

To simulate changes in interaction properties (e.g., compliant surfaces or softened fingertips), frequency-domain subtraction was applied to construct compensation signals. Specifically:

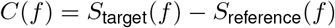

where *S*_target_(*f* ) is the FFT of the harder reference material (e.g., wood or metal) and *S*_reference_(*f* ) is the FFT of the baseline condition (e.g., foam or bubble-modified finger). The inverse FFT of *C*(*f* ) was computed to generate time-domain compensation signals. This approach effectively overlays the missing spectral components of a harder material onto the degraded baseline condition.

#### Signal Normalization and Upsampling

All signals were RMS-normalized to a target amplitude of 0.35 G to ensure perceptual comparability across materials and signal-generation methods. Signals were then upsampled from the acquisition rate (2000 Hz) to the playback rate (44.1 kHz) using a windowed-sinc interpolation filter. Final signals were saved in 16-bit mono WAV format for playback.

#### Real-Time Signal Generation

We used three approaches to synthesize material-specific vibration cues:

- **Data-driven Model** This approach captured real acceleration signals from finger-material interactions and replayed them directly, offering high realism (*4, 20, 21*).
- **Decaying Sinusoidal Model** Signals were synthesized as

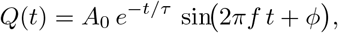

where *A*_0_ is the initial amplitude, *τ* the decay time constant, *f* the carrier frequency, and *ϕ* the phase. To minimize onset/offset artifacts, each signal was multiplied by a Gaussian window (∼50 ms total duration). Parameters (*τ, f* ) were chosen to approximate recorded spectra for each material (e.g., foam: *τ* = 5 ms, *f* = 85 Hz; metal: *τ* = 20 ms, *f* = 400 Hz).
**Ricker Wavelet Model** The Ricker wavelet was defined as:

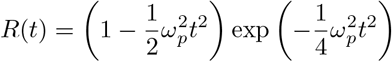

Peak frequencies *ω*_*p*_ were chosen to approximate dominant spectral components of each material ( (*33, 34*)). The duration of each waveform was fixed at 50 ms with zero-padding.

All signals were within trigger-to-output latency of *<* 1 ms of contact detection (measured with oscilloscope).

#### Playback and Actuation

Photoelectric sensors positioned above the tapping surface detected finger contact events. Sensor triggers initiated signal playback on a Bela board running Xenomai real-time Linux. Signals were routed through a TPA3116D2 Class D amplifier and delivered to Lofelt L5 actuators affixed to participants’ fingernails. This ensured both low-latency delivery and high-fidelity reproduction of vibration cues. Signals were band-limited to 30-1000 Hz (8th-order zero-phase Butterworth). Peak acceleration did not exceed 1.2 G and remained below published discomfort thresholds for fingertip vibrotactile stimulation. Participants could request breaks at any time; no adverse events were reported.

## Experimental Procedures

Participants’ index fingers were lightly wrapped with tape to reduce confounding cues (e.g., temperature, skin texture). All studies used the same actuation and triggering pipeline (latency *<* 1 ms) and were run single-blind with randomized trial order. Participants were instructed to use brief, light taps at a comfortable and repeatable pace (“as if quickly checking a surface”). The participants were then given a demonstration where the finger were fully raised without raising the hand at a pace of approximately 90 BPM. We selected tapping to constrain the exploratory mode and reduce variability associated with sustained pressing, while allowing the rendered transient to be coupled consistently to contact onset. We matched RMS acceleration per stimulus (equivalently, total spectral energy integrated over 30-1000 Hz via Parseval’s theorem) and refer to this matched quantity as *total spectral energy (RMS-matched*).^1^

### Fingertip modification experiments (Experiments 1-2)

#### Experiment 1: Hardness discrimination with softened fingertip (2AFC)

- **Goal**. Test whether added spectral energy restores relative hardness judgments when fingertip stiffness is reduced.
- **Participants & setup**. Ten participants wore an inflatable bubble actuator on each index fingerpad (5 psi). Both hands tapped identical rigid wood blocks; one block delivered a vibration transient at contact, the other was silent.
- **Stimuli**. A 3×5 set: three waveform types (data-driven, decaying sinusoid, Ricker wavelet) crossed with five RMS-matched energy levels (No-signal control; foam-, silicone-, rubber-, wood-/metal-like total energy). Signals were brief (*∼*50 ms), aligned to contact onset.
- **Task & design**. 2AFC (“Which tap felt harder, left or right?”). For each waveform, all ordered pairs of the five levels that differed in energy were tested once per hand (20 pairwise comparisons per waveform), then repeated once (2 reps/hand). Total per participant: 60 comparisons, randomized. Session duration: 30-45 min.
- **Planned analysis**. Accuracy per contrast; binomial tests versus chance for adjacent pairs; repeated-measures ANOVA on accuracy with factors `difficulty` (adjacent pair) and `waveform`; confusion matrices for descriptive summaries.

#### Experiment 2: Material matching with softened fingertip (5-AFC)

- **Goal**. Test whether spectral energy alone conveys material identity when fingertip mechanics are softened.
- **Participants & setup**. Ten participants, bubble actuator on the non-dominant index finger (taps a wood block with vibration); dominant index finger explores five real reference materials (foam, silicone, rubber, wood, metal) without vibration.
- **Stimuli**. Fifteen signals (3 waveforms *×* 5 RMS-matched energy levels), one transient per trial.
- **Task & design**. 5-alternative forced choice (“Which real material best matches the rendered vibration?”). Each of the 15 signals presented 5 times (75 trials), randomized, after one practice trial. Duration: 30-60 min.
- **Planned analysis**. Confusion matrix; ordinal (Spearman) correlation between energy level and chosen material rank; mutual information; two-way ANOVA (factors: `energy level, waveform`) on response rank; Tukey post hocs for adjacent levels.

#### Surface modification experiments (Experiments 3-4)

##### Experiment 3: Hardness discrimination on a soft surface (2AFC)

- **Goal**. Test whether added spectral energy can override a uniformly soft substrate in relative hardness judgments.
- **Participants & setup**. Ten participants, bare fingertips, two identical foam blocks; one block delivered vibration, the other was silent.
**Stimuli**. Same 3 × 5 set as Experiment 1 (three waveforms × five RMS-matched energy levels), transient aligned to contact.
- **Task & design**. 2AFC (“Which tap felt harder, left or right?”). Total: 60 pairwise comparisons per participant (as in Exp. 1), randomized. Duration: 30-45 min.
- **Planned analysis**. As in Exp. 1 (binomial tests for adjacent contrasts; rmANOVA on accuracy with `difficulty` and `waveform`; confusion matrices).

##### Experiment 4: Material matching on a soft surface (5-AFC)

- **Goal**. Test whether spectral energy conveys material identity when the touched surface is compliant.
- **Participants & setup**. Ten participants, bare fingertip taps a foam block with vibration; dominant hand explores the five reference materials.
**Stimuli**. Twelve signals (3 waveforms × 4 RMS-matched energy levels spanning foam→ metal; highest two levels can be collapsed if needed based on calibration range).
- **Task & design**. 5-alternative forced choice. Each signal repeated 5 times (60 trials), randomized, after one practice trial. Duration: 30-60 min.
- **Planned analysis**. As in Exp. 2 (confusion matrix, ordinal correlation, mutual information, two-way ANOVA with `energy level` and `waveform`).

##### Experiment 5: Cue-conflict (surface compensation) paradigm

- **Goal**. To test whether perceived material identity is dominated by total spectral energy (RMS) or by dominant frequency, we dissociated these cues in rendered vibrations delivered while tapping a soft substrate.
- **Participants and setup**. Ten participants tapped a physical foam block with a bare finger. Vibrotactile playback was delivered to the fingernail via the Lofelt L5 actuator (latency *<* 1 ms), as in prior experiments.
**Stimulus construction**. We created a 3×3 factorial set by crossing three RMS targets with three dominant-frequency classes, yielding 9 stimuli (3 congruent, 6 incongruent):
  - **RMS (“energy”) targets:** values matched to the band-limited (30-1000 Hz) RMS acceleration measured from our data-driven recordings for *foam, rubber*, and *wood*. RMS was computed in the frequency domain and verified in time domain (Parseval’s theorem).
  - **Dominant-frequency classes:** peak frequencies characteristic of the same three materials, grouped as ≈ 85 Hz (foam-like), ≈ 180 Hz (rubber-like), and 350-400 Hz (wood-like).

Each stimulus was an exponentially decaying sinusoid, windowed to a brief transient and aligned to contact onset:

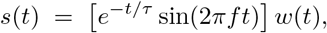

with *f* ∈ {85, 180, 350-400} Hz, *w*(*t*) a Gaussian/Hann onset window (total duration ∼50 ms), and *τ* chosen to match the decay constant of recorded transients. After generating *s*(*t*) at the desired *f*, we scaled the *time-domain* amplitude so that the playback signal’s band-limited RMS (30-1000 Hz) matched the assigned energy target. We refer to this matched quantity as *total spectral energy (RMS-matched*).

*Congruent* conditions paired foam-RMS with 85 Hz, rubber-RMS with 180 Hz, and wood-RMS with 350-400 Hz. *Incongruent* conditions crossed each RMS with the two non-matching frequency classes.

- **Task and design**. On each trial, participants tapped the foam surface once and received one of the nine stimuli at contact. They then made a 3-alternative forced choice (3AFC): *foam, rubber*, or *wood*. Each of the 9 conditions was presented 5 times (45 trials total, 10-15 minutes), in randomized order, following one practice trial. Sessions were conducted single-blind.
- **Planned analysis**. Primary outcome was the chosen category per trial. We fit a multinomial (or separate binomial) mixed-effects logistic regression with fixed effects of RMS target and frequency class (and their interaction), and random intercepts for participant. Confusion matrices and energy-rank correlations were computed as descriptive summaries.

### Calibration and Control

#### Energy matching

We matched RMS acceleration per stimulus (equivalently, the total spectral energy integrated over 30-1000 Hz by Parseval’s theorem). We refer to this quantity as “total spectral energy (RMS-matched).” Durations and windowing were held constant across waveforms.

#### Amplitude Calibration

RMS amplitude was standardized across materials and methods to ensure that perceived intensity did not confound perceived hardness. A pilot perceptual matching task (n = 4) confirmed that amplitude-normalized waveforms were not systematically perceived as more or less intense. Calibration values are provided in Table S2.

#### Control Signals

Three “null” signals were included for each generation method:

1. A flat-zero waveform (no vibration)
2. A low-amplitude noise burst (sub-threshold)
3. A neutral sine pulse (non-material specific)

These were used to evaluate baseline response rates and exclude participants with response bias.

### Trial randomization and data collection

Each trial began when the participant tapped the assigned contact surface (foam or wood) with a prepared index finger. Upon sensor trigger, a vibration cue was delivered within *<* 1 ms. Participants were free to alternate between stimuli (left/right or physical/rendered) before responding via a button box.

Stimulus order was randomized *within participant*. For each block, left/right assignment, waveform type, and energy level were independently randomized *without replacement*, ensuring equal counts per condition. Block order (discrimination vs. matching) was counterbalanced across participants using a balanced Latin square. Sessions were conducted single-blind: participants were unaware of condition and the experimenter did not see trial assignments during data collection.

One limitation to the experimental design is that impact force and velocity were not instrumented or force-controlled during the psychophysical tasks, with an attempt to minimise variation only by given guidance on height and rate of tap to the participants. Therefore, natural trial-to-trial variability in impact mechanics may have contributed noise to the sensory signals. However, contact onset was detected independently of impact magnitude and triggered the controlled vibration stimulus within 5 ms. Because the same instructed tapping procedure and randomized design were used across conditions, and because responses showed systematic monotonic effects of the experimentally controlled energy levels across participants and waveform families, such variability is unlikely to account for the principal effects.

#### Data Collection Interface

Trials were managed via a custom Python interface using PyDAQmx and PyQt5, with real-time communication to the Bela board over UDP. Participant responses and timing logs were recorded with 1 ms resolution.

### Additional Notes and Figures

- **Figures:**
  - Figure 9a: Fingertip discrimination setup
  - Figure 9b: Fingertip matching task
  - Figure 9c: Surface discrimination task
  - Figure 9d: Surface matching task
  - Additional setup and equipment photos: Figures 10a, 10b, 10c
- Foam was chosen as the compensation surface due to its low hardness and consistent tactile properties.
- The actuator on the dominant hand during matching tasks served only as a mass balancer.

**Figure 9.**
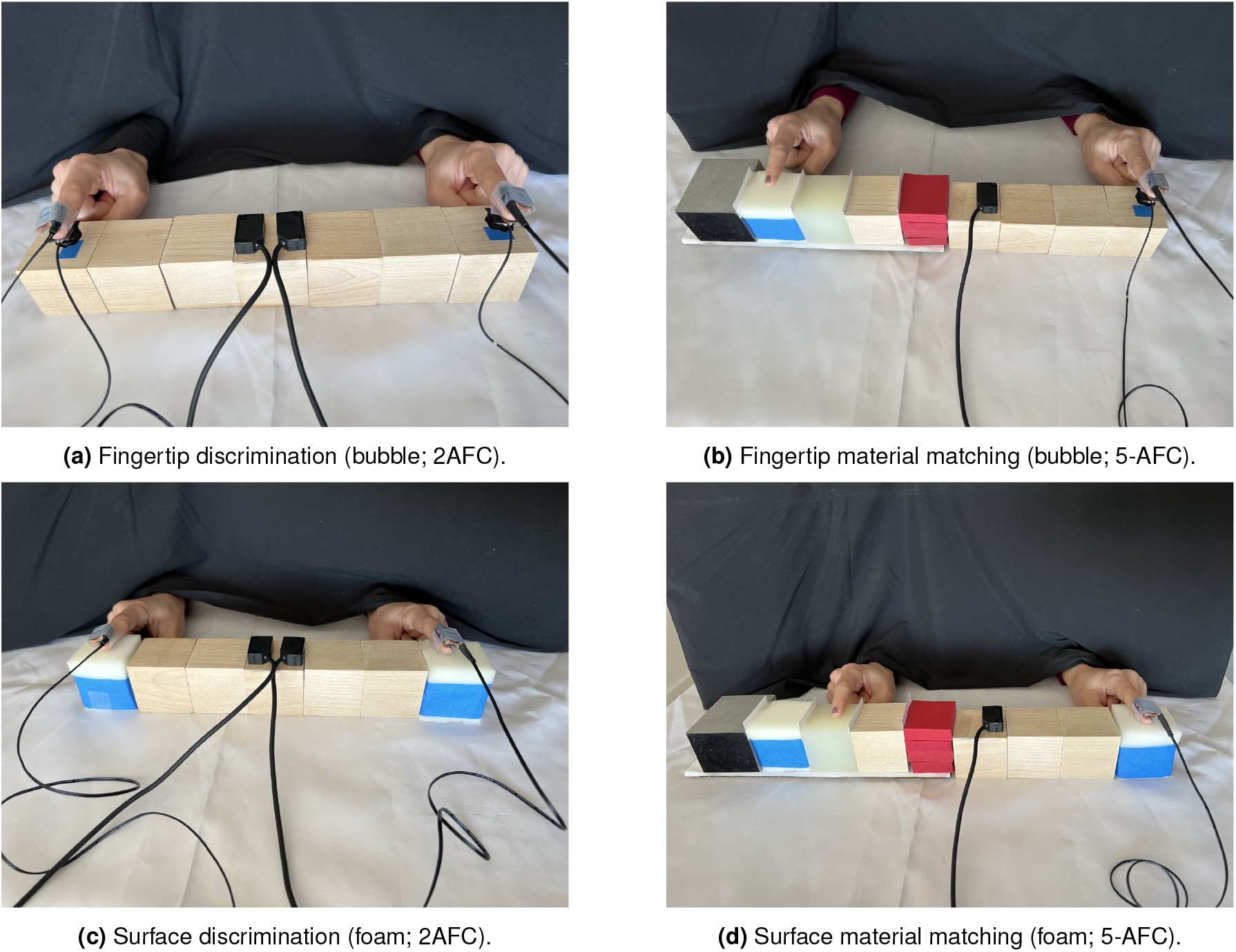
Behavioral paradigms and tasks. **A** Fingertip-discrimination setup: bubble-softened index fingers tap two rigid blocks; one tap includes vibration (2AFC: harder). **B** Fingertip-matching setup: bubble-softened finger taps wood with vibration and is matched to a real material (5-AFC). **C** Surface-discrimination setup: bare finger taps two foam blocks; one tap includes vibration (2AFC). **D** Surface-matching setup: bare finger taps foam with vibration and is matched to a real material (5-AFC). Triggers align transients to contact with sub-millisecond latency.

**Figure 10.**
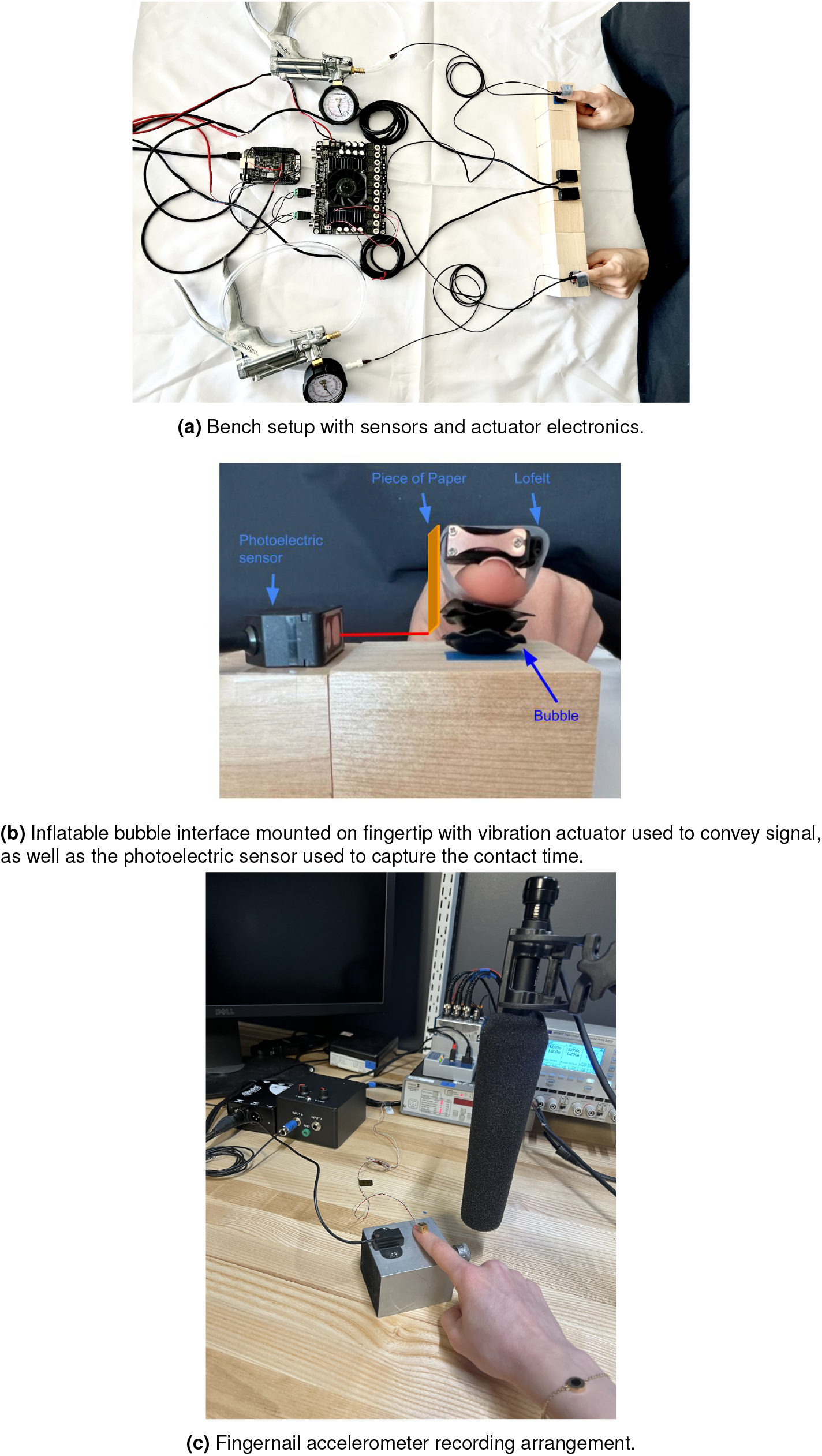
Apparatus and instrumentation. **A** Experimental bench with photoelectric contact sensors and Bela real-time playback. **B** Inflatable fingertip bubble (5 psi) used to reduce fingertip stiffness while leaving the nail accessible for actuation. **C** Recording setup for data-driven transients: tri-axial accelerometer mounted to the fingernail captures contact-induced vibrations for later synthesis and energy matching.

## Footnotes

1 RMS matching procedure and values are detailed in Supplementary Methods, Signal Calibration.

